# Spatial structure and pathogen epidemics: the influence of management and stochasticity in agroecosystems

**DOI:** 10.1101/2020.06.19.161810

**Authors:** Zachary Hajian-Forooshani, John Vandermeer

## Abstract

1. Organisms susceptible to disease, from humans to crops, inevitably have spatial geometry that influence disease dynamics. Understanding how spatial structure emerges through time in ecological systems and how that structure influences disease dynamics is of practical importance for natural and human management systems. Here we use the annual crop, coffee, *Coffea arabica*, along with its pathogen, the coffee leaf rust, *Haemelia vastatrix*, as a model system to understand how spatial structure is created in agroecosystems and its subsequent influence on the dynamics of the system.
2. Here, we create a simple null model of the socio-ecological process of death and stochastic replanting of coffee plants on a plot. We then use spatial networks to quantify the spatial structures and make comparisons of our stochastic null model to empirically observed spatial distributions of coffee. We then present a simple model of pathogen spread on spatial networks across a range of spatial geometries emerging from our null model and show how both local and regional management of agroecosystems interact with space and time to alter disease dynamics.
3. Our results suggest that our null model of evolving spatial structure can capture many critical features of how the spatial arrangement of plants changes through time in coffee agroecosystems. Additionally, we find that small changes in management practices that influence the scale of pathogen transmission, such as shade tree removal, can result in a rapid transition to epidemics with lattice-like spatial arrangements but not irregular planting geometries.
4. The results presented here may have practical implications for farmers in Latin America who are in the process of replanting and overhauling management of their coffee farms in response to a coffee leaf rust epidemic in 2013. We suggest that shade reduction in conjunction with more lattice-like planting schemes may result in coffee being more prone to epidemic-like dynamics of the coffee leaf rust in the future.

## Introduction

Organisms susceptible to disease, from humans to crops, inevitably have spatial geometry that influences disease dynamics. While it may be argued that spatial components of disease-host systems in mixed environments are less important (e.g. plankton), it is certainly true that most plants and animals have non-trivial spatial structure, whether exogenously imposed by abiotic environment (Gratzer et al. 2004) or emerging endogenously from ecological dynamics (Li et al 2016). It has been a standard epidemiological question to ask how disease propagates through space (Keeling et al. 1999; Park et al. 2002; Balcan et al. 2009; Craft et al. 2010), but less obvious is how the space is constructed in the first place and how that space influences subsequent disease dynamics. At one extreme, a feedback likely exists between host and disease, where hosts may alter their spatial distribution in response to the presence of disease, such systems may include humans (Levine & Levine 1994). On the other hand, there exist many hosts-pathogen systems where hosts are unable to alter their spatial distribution over the course of pathogen dynamics, such as crops. Here we focus on the latter case and use crops as a model system to explore the generation of spatial structure and how subsequent spatial structure influences pathogen dynamics.

Likely all crops in cultivation are susceptible to pathogens, and historical examples such as The Great Famine in Ireland and the future threats such as Panama disease on bananas highlight the implications for understanding how pathogens operate within agroecosystems. Although the construction of spatial structure is often a direct consequence of agricultural intensification, its impact on pathogen dynamics in agriculture is often overlooked by agronomists. We argue that the construction of space becomes central to the understanding of the control dynamics of the system, especially in the case of agricultural pathogens. Here we use the agent of the coffee rust disease, *Haemelia vastatrix*, where the spatial scale of coffee plants is evidently important with regard to disease spread (Avelino et al. 2012; Vandermeer et al., 2017; Vandermeer & Rohani 2014). Disease transmission is from plant to plant on any given farm, but from farm to farm and region to region, at larger scales, and the choice of a framework on which to build theoretical understanding of this system is conditioned partly by its evident spatial context.

The local transition component, i.e., from coffee tree to coffee tree is clearly dependent on the spatial arrangement of the trees. However, that spatial arrangement is a consequence of farmer decisions about initial planting with the important addition that the spatial pattern is modified by continual replanting in spaces where individual trees had become damaged or died. It is evident that the planting design is an evolving lattice, beginning with ordered rows and interplant distances, but evolving over time, with the dynamics of replanting, a response to thinning, from a variety of causes, including the coffee rust disease itself. Consequently, the pattern of disease occurrence in this system is conditioned first by the structure of the coffee plant distributions (effectively a socioecological process) and second by the dynamics of transmission (mainly an ecological process). We find it convenient to represent this phenomenon as a network model, with dynamic forces creating the network structure. For the coffee plants themselves, it is a combination of intentional imposition of a regular lattice (planting coffee bushes in well-defined rows) coupled with the pruning and natural deaths requiring replanting on a periodic basis. Subsequently, the rust disease dynamics unfold on this dynamically constructed network, resulting in a new network, in its most elementary form with the presence or absence of the rust. Thus, we have a meta-network, in which certain dynamic forces (planting, pruning and replanting coffee plants) create a particular network structure on which the dynamic forces of disease transmission create a network pattern of the coffee rust disease itself.

Here we employ a straight-forward approach to explore the interplay between the generation of underlying spatial geometries of hosts and the implications of those geometries for pathogen dynamics. First, we develop a simple null-model to simulate the socio-ecological processes that determine the underlying geometries of plants within a coffee agroecosystem (although this framework will be applicable to any agroforestry system). We then use a spatial network approach, as employed by others (Keitt et al. 1997), to determine the types of spatial geometries our null-model produces and how they compare to the empirical geometries observed in actual coffee agroecosystems. Finally, we simulate pathogen spread on the range of spatial geometries generated by our null-model and study the implications of the interplay between the management of these systems with the spatial geometries driven by stochasticity.

## Methods

### Null model of evolving plant spatial geometry

Despite the fact that coffee bushes are often planted with the intention of a strict lattice structure (planted in rows), the real distribution of coffee plants on a farm rarely reflects perfectly that initial intent. As time goes by, some coffee bushes die and usually are replanted, but rarely in precisely the same location, leading eventually to a loss of the initial planting pattern. While to the farm worker these small deviations may not seem consequential for the dynamics of pathogens and pests, her we explore how these deviations accumulate over time and what their impacts are on pathogen dynamics. Although a host of complicated local factors are involved in planting decisions, we initially approach the problem with a null model of planting spatial evolution. We begin with plants arranged in a lattice bound within a *x* and *y* coordinate range and modify the structure over time. The simple model simulations stochastic death and replanting within an area within the proximity of the prior plant. The model is:

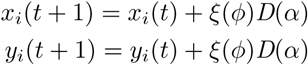

Where

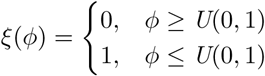

and *x*_*i*_(*t*) and *y*_*i*_(*t*) represent the two coordinates corresponding the position of plant *i*, at time *t, ϕ* is the mortality probability for plant *i, U(a,b)* is a uniformly distributed random variable with range 0,1, *ξ*(*ϕ*) is the death/replanting rate, and *D*(*α*) is a random variable with mean *a*, stipulating the “replanting radius”. When *ξ*(*ϕ*) = 1 the x and y coordinates are modified by *D*(*α*). The simulations were run iteratively for each plant in the plot 200 times.

To understand how our null model approximates the planting arrangements of real agroecosystems, we use an empirical data set of three 20 × 20 m plots on an organic coffee farm in the Soconusco region of Chiapas, Mexico. Given these three 20×20m plots with differing numbers of plants, for each comparison, the simulated evolution of spatial structure was done with the same planting density as the real plot it was intended to simulate. By controlling for the planting density, our null model of plot spatial evolution allows us to make comparisons with our empirical 20×20 plots and understand to what extent our null model approximates the actual spatial geometry.

### Quantification of spatial structure

In the same vein as our prior work on this system, which sought to understand the spatial scale of pathogen transmission (Vandermeer et al. 2018), we argue that the sub-graphs (or connected components) provide a useful way study the local spread of pathogens in space. To quantify the spatial structure for both empirical and model generated plots, we borrow from the common framework of graph theory to understand spatial structure (Fletcher et al. 2016).

Our basic framework employs a threshold radius or critical distance (*D*_*crit*_) which defines edges between points in space, where all nodes are connected by edges equal or smaller than *D*_*crit*_, thus forming a sub-graph within the space. For our purposes here, the number of sub-graphs within a coordinate system is useful in that the same measure has a biological interpretation when modeling the pathogen in space. For example, it gives us information about the minimum number of outside infections needed to fully infect a plot, allowing for the separation of intra and inter sub-graph dynamics. The use of sub-graphs to describe the spatial structure is also quite useful in that that we are working with relatively small networks (*<* 400 nodes), and spatial scales, *D*_*crit*_, where the spatial network is not fully connected.

To understand the extent to which the simulations approximate the empirical patterns, we plot Δ_*s*_, the difference in the number of sub-graphs in the actual structure minus the number of sub-graphs in the simulated structures. For a perfect spatial approximation in terms of number of subgraphs we expect a Δ_*s*_ = 0. For each simulation, we extracted the pattern at the first step and subsequently every 10 steps through 100 rounds of replanting.

### Model of Spread on Spatial Networks

When considering the dynamics of a pathogen unfolding on spatially structured networks of host plants, *D*_*crit*_ takes on an important biological interpretation. It is effectively the maximum radius the pathogen is able to spread to neighboring plants, the scale of transmission. Similar to the quantification of spatial structure in the previous section, *D*_*crit*_ is used to generate different spatial network structures that reflect an inevitable distribution of the pathogen in space via a simple spreading processing to neighbors. The number of sub-graphs give us biologically relevant information for the dynamics of the pathogen that the use of a dispersal kernel does not highlight outright. As mentioned above, a give distribution of sub-graphs for a *D*_*crit*_ gives us the minimum number of outside infections needed to infect a total are in cultivation. It has been shown that the intensity of pathogen infection is correlated within a sub-graph empirically (Vandermeer et al. 2018), and here we use this as a simplifying assumption in our model. We constructed a simple model of pathogen transmission to explore the consequences of the different patterns that emerge from the null model of pattern formation, based on our previous model of this system (Vandermeer et al. 2017).

We start with a given number of plants or nodes within the system, *P*.

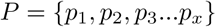

Where *x* is the number of plants in the system.

The model is initialized with a coordinate system which corresponds to the spatial position of each plant, which emerges from the simulations from the plot spatial evolution. Then a spatial scale of pathogen transmission is stipulated via a *D*_*crit*_ which in turn creates a collection of spatial sub-networks (frequently referred to as “connected components”), called *C*

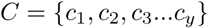

Where *y* is the number of sub-graphs in the system. Note that each subnetwork in *C* contains a unique collection of plants from *P* corresponding to a given scale of pathogen transmission *D*_*crit*_. Here is an illustrative example:

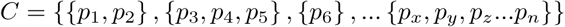

Note that the indices for each plant, *p*, are unique across all subsets within *C*.

In the model, we keep track of all the infected plants with *I*, which is initialized as empty set.

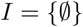

For each time step in the model we iterate through all nodes (plants) in *P*, and there is a fixed probability,*β*, that a given plant becomes infected. If *p*_*i*_ (the *i*th plant in *P*) becomes infected via

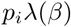

Where

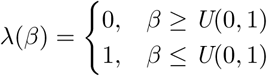

then the cluster,*C*_*j*_, which is a subset of *C* that contains *p*_*i*_, is joint in union with *I*. This is done for all *p*’s where *λ*(*β*) = 1, in other words, where there was a successful infection.

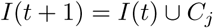

In summary, the basic dynamic of the model is: the set of infected plants, *I*, at time *t* + 1 is equal to the set *I* at time *t* joint in union with the cluster (*C*_*j*_), which is a subset of *C*, that contains all of the plants, *p*_*i*_, that were infected successfully via *λ*(*β*) = 1. For each times step we are primarily interested in the number of infected plants which can be represented as the cardinality of *I*,

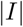

Conceptually, note that each sub-network represents the extent to which the pathogen instantaneously spreads from a single infected plant to all plants in that sub-graph. We use the inevitability of the spread within a sub-graph as a simplifying assumption and assume that all hosts that fall within the sub-graph denoted by the scale of the spread, *D*_*crit*_, become infected instantaneously. Given that prior work has shown that infection rates tend to be more similar within a sub-graph than between them (Vandermeer et al. 2018), this abstraction simplifies the system and provides for a focus on the interplay of pathogen dynamics and the spatial geometry at the scale of the plant. With the assumption of instantaneous spread within a sub-graph, our model only has one parameter associated with the epidemic process, the probability of a random plant in the plot being infected, *β*, which can be thought of as being a measure of the regional pathogen propagule density.

## Results

### Plot evolution and approximations of empirical structure

Empirical spatial distributions correspond qualitatively to the distributions predicted from the null model (Figure. 1). Considering the pattern of subgraph emergence as a function of *D*_*crit*_, we expect that as time advances (iterations in the model) the model will approximate the empirical (as seems to be the case in Figure 1). Data for the three empirical plots are roughly approximated by the null model for various spatial scales (values of *D*_*crit*_). The range of shades in Figure 2 show the variation in plot evolution, where light grey is the lattice and dark red is after 100 rounds of replanting. The simulations start far from the empirical distributions and move towards them (i.e., Δ_*s*_ = 0).

**Figure 1:**
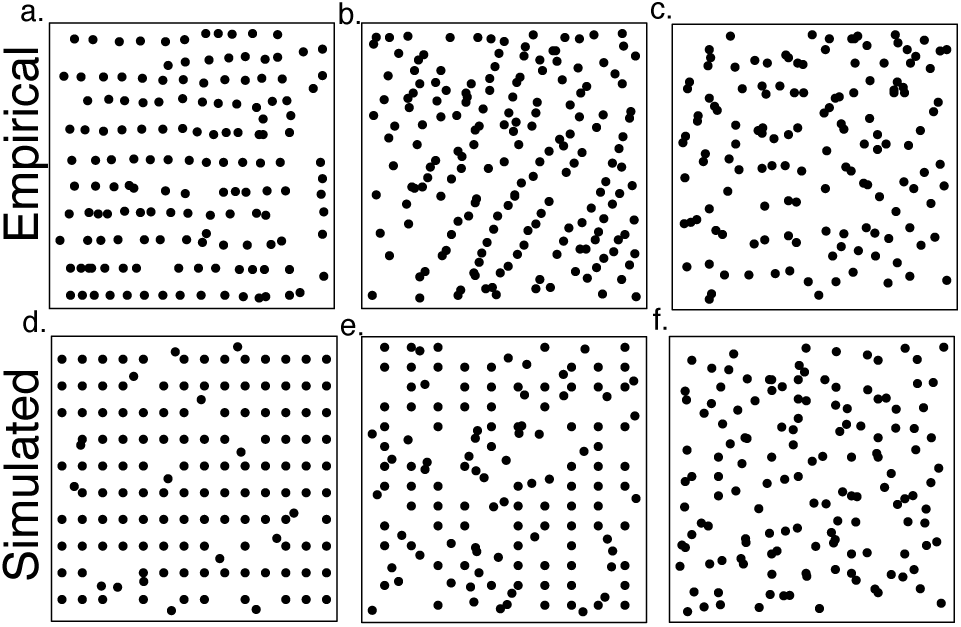
Three 20 x 20M plots illustrating the position of all coffee plants. a. a one-year old plot, b. a 4 year old plot, c. approximately a 15 year old plot d, simulated plot after 50 time units, e. simulated plot after 100 time units, f. simulated plot after 500 time units.

**Figure 2:**
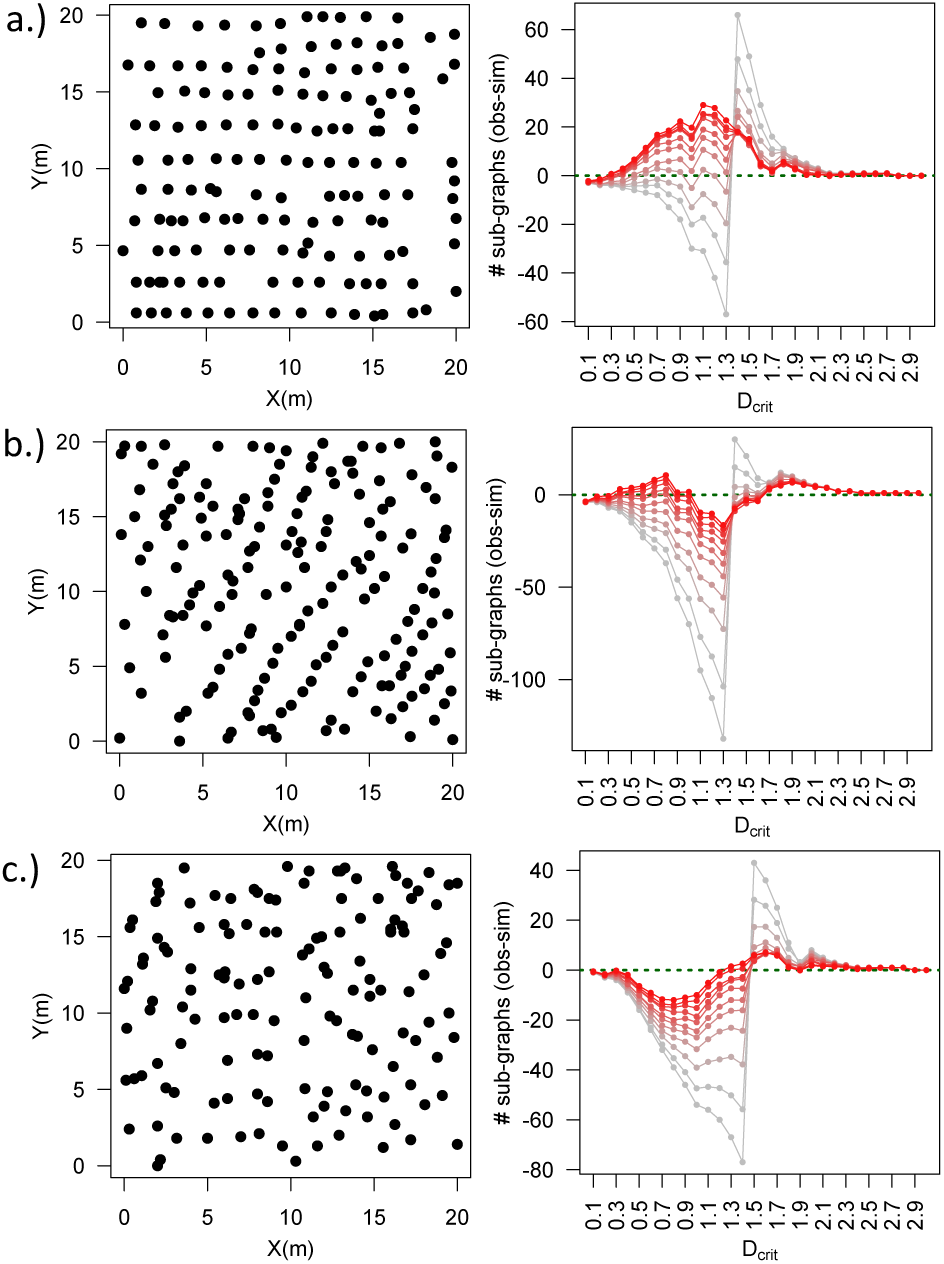
(a-c) Δ_s_ as a function of the critical distance (parameter a in equation set 1), at various stages in the evolution of plot structure using the null model. Shading goes from light (the first stage in the simulation) to dark red (final step of simulation). Note that the green dashed line corresponds to a 1:1 approximation of the model to the empirical plots. The model clearly approaches the null expectation as the simulation proceeds.

The largest deviations (Δ_*s*_) are typically found at the distance that separates rows of the lattice. This suggests that the empirical planting geometries are more clustered and over dispersed at scales that the model cannot approximate. For example, the empirical plot in Figure 2a. shows the model consistently unable to approximate at *D*_*crit*_ from 1.3-2m, and we see in the empirical data that this likely emerges from irregularities within row structure. It is evident from Figure 2a. that the deviation from the lattice emerges from missing plants and clustered plants but within the row structure itself. While simulations move plants away from the lattice structure randomly, the empirical data suggest that attempts to maintain semblance of row structure results in plants being replanting within the row but in an over dispersed or clustered fashion. Similar deviations are found between the empirical plots and simulations in Figure 2b and c and are consistent prior to the scale that join rows of the lattice, as denoted by the grey line from the simulations. These deviations occur because the simulated plots are more clustered at these smaller distances as shown by the approximations being below the zero line.

### Modeling pathogen spread on spatial networks

Using time to reach 90% infection as a state variable, we illustrate its response to the two variables of interest, “plot evolution time” which is to say the time the null model is permitted to run, and the *D*_*crit*_ parameter of the connect component determination (the model as discussed in the methods section above). In Figure 3 we summarize the general dynamics of the system from a two-dimensional parameter sweep of, 1) the scale of the pathogen transmission (*D*_*crit*_), 2) the time steps involved in the plot evolution simulation, and 3) the state variable, time to epidemic (time to reach 90% of the trees infected). A pathogen spreading across the different plot geometries (evolution time) reaches epidemic status regardless of spatial geometries due to limited transmission to neighboring plants the majority of which fall outside of the threshold (*D*_*crit*_). At low transmission levels it is apparent that plants become more clustered as the plot geometry evolves away from a lattice-like structure resulting in a small but detectable difference in the time to epidemic. A similar feature is observed at large scales of pathogen transmission due to almost the entire plot being connected resulting in any plant becoming instantaneously infected once a colonizing infection reaches the plot.

**Figure 3:**
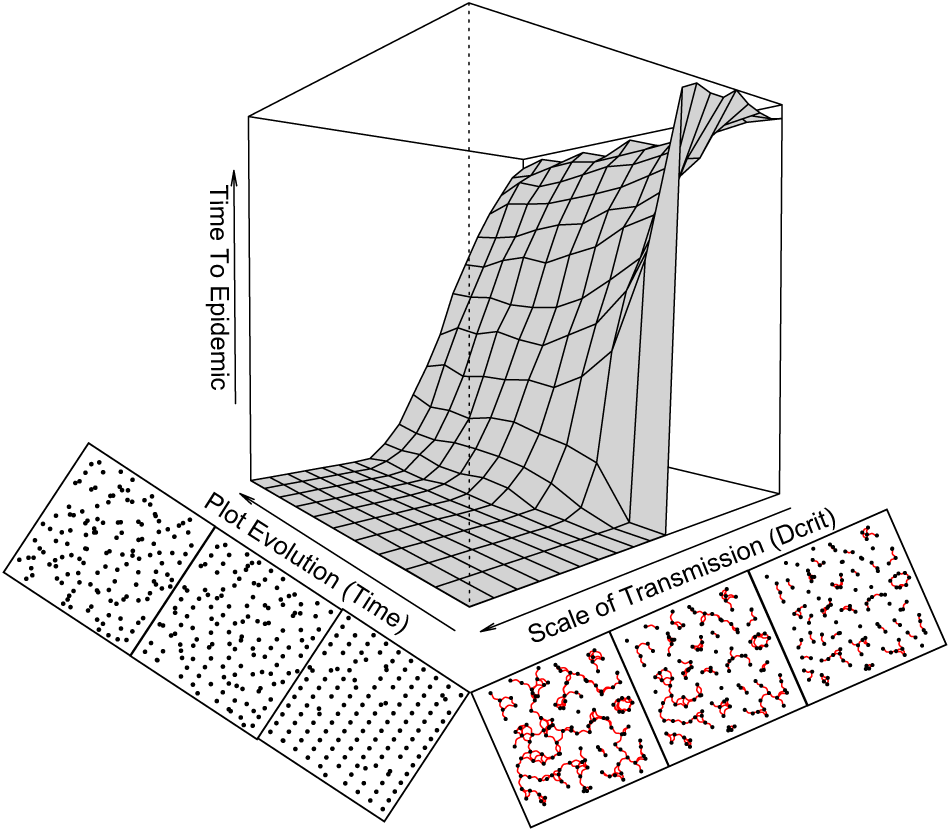
Shows the time until 90% infection across a range of scales of pathogen transmission (D_crit_) as well as planting geometries generated by the plot evolution model. Plots along the two axes are to illustrate the changing spatial network structure along scale of spread and the changing panting geometry in the plot evolution model. Note that the regional infection probability here is at 0.1

It is intuitive that at very large transmission values, the pathogen will move quickly to infect the whole plot and at smaller scales it will move slowly, regardless of the planting geometry. It is at intermediate transmission levels that we find non-obvious interactions with the geometry of host plants. For these intermediate scales we find that lattice-like geometries are sensitive to increases in transmission and generate a drastic jump in pathogen dynamics where the time to epidemic shows a pattern similar to that of a critical transition (Figure 3.). As the plot becomes less lattice-like through stochastic host death and replanting this jump in dynamics becomes less pronounced. At the two extremes of host planting geometries, we see critical transition-like behavior for highly organized lattice-like arrangements and a gradual chance for more unorganized pseudorandom arrangements. These results suggest that a more random-like pattern of the host buffers drastic changes in the overall dynamics of the pathogen.

Given the basic biology of most pathogens, it makes sense to think about not only the dynamics within a plot but also how the regional dynamics impact the system. Our model results suggest that the dynamics of the spatial host-pathogen system qualitatively changes as the probability of plants being infected from outside of the plot increases (Figure 4). As the regional infection probability increases, the interaction between the spatial geometries of the hosts and level of pathogen transmission becomes less pronounced. The critical transition-like behavior observed for relatively low regional infection probabilities is buffered as the regional infection probability increases, suggesting that under epidemic levels of a pathogen in the environment, the spatial arrangement of hosts on a given farm becomes less important for the overall dynamics of the system.

**Figure 4:**
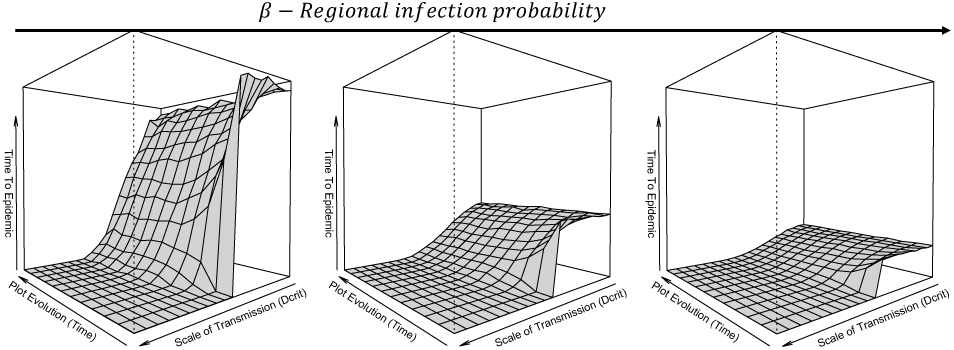
Shows how the pathogen dynamics change as the regional infection probability increases. The first figure is β = 0.1, then β = 0.2, then β = 0.5.

## Discussion

The actual management practices that create the spatial geometries of plants in agroecosystems likely emerge from socio-ecological processes structured by a host of influences, from cultural practices to the socioeconomic status of the farmer. Our approach here has been to try and recreate the range of observed spatial planting geometries by using a simple null model that strips away most of these real-world complexities. We show that a simple process of stochastic plant death and replanting within a small radius surrounding the dead plant can recreate many of the features observed in the real distribution of planting geometry. Furthermore, we suggest that the observed spatial geometries in agroecosystem can be the result of different snapshots in time of those same dynamic process (Figure 1). The comparisons between our model and empirical data (Figure 2) provides support for the idea that at least for the sampled plots, the distributions of plants fall along different times of this stochastic death and replanting process. For the most lattice-like geometry of our empirical plot (Figure 2a.) the model passes through the 1:1 approximation of our empirical data for a wide range of spatial scales, suggesting that early stages in the simulations that move away from the lattice approximate it better than later more unorganized steps in the model. Furthermore, our simulations pass that same 1:1 approximation but at a much later time in the simulations for our plot of intermediate lattice structure (Figure 2.b), while our plot farthest from the lattice structure (Figure 2.c) is never well approximated by our model.

The interaction of spatial structure and spatial transmission of the pathogen suggests that they interact in a non-linear way. We observe that there is a critical transition-like behavior that emerges from the interaction of both the scale of pathogen transmission and the underlying spatial geometry of the system. Relatively small changes in pathogen transmission (*D*_*crit*_) with lattice-like spatial geometry can lead to a dramatic jump in dynamics of the pathogen (Figure 3). Thus, with lattice-like planting, the pathogen may be held at relatively low densities, but a small change in management that may influence of scale of pathogen spread (discussed below) can result in a devastating shift in dynamics. The uniform nature of the lattice creates the conditions that, once the threshold that connects rows is met, the whole plot becomes a connected network on which for the pathogen can spread across. As the death and replanting process that moves the spatial pattern away from a lattice planting geometry, it disrupts row structures and subsequently buffers the critical transition-like jump in pathogen dynamics. Thus it might be expected that uniform lattice-like planting geometries are far more sensitive to small changes in the scale of transmission of the pathogen and even a small amount of disruption from the highly organized state can buffer against variability in pathogen spread. As we move away from the lattice-like geometry, there may be higher pathogen infection at low transmission due to some clusters of plants, but that same geometry prevents the pathogen from spreading through the whole area. Furthermore, as the regional infection probability increases the qualitative results of the model stay the same but they are damped, and the critical transition-like behavior is almost non-existent for high regional infection probabilities. This suggests that under regional epidemic regimes when enviornmental pathogen loads are so high, that the host geometry makes little impact in buffering the pathogen dynamics.

The results presented here have practical significance for the management of pathogens in agricultural systems, and in particular for the system which inspired the study, the coffee leaf rust (CLR). The CRL propagates both as a random propagule rain, spores arriving from the regional pool of spores in the environment, and from plant to plant on a local level through local wind current instabilities, branch-branch-contact, and splashing (Vandermeer et al. 2017, Avelino et al 2015). Our model simulations mainly focused on the local level transmission and how that interacts with the planting geometry of the agroecosystem. There are a number of management factors within coffee agroecosystems that have the potential to influence parameters associated with the scale of the CLR transmission (Avelino et al. 2004). Shade is one of the most commonly managed aspects of coffee agroecosystems and its impact on the dynamics of the CLR has been contentious with some reporting beneficial impacts of shade reducing CLR (Soto-Pinto et al, 2002), and others reporting the opposite (Lopez-Bravo et al. 2012). The classic recommendation has been to reduce shade to manage the CLR, as the microclimatic implications of shade such as increasing humidity could potentially be beneficial to the germination of spores (Staver et al. 2001). Bourot et al. (2016) provided evidence that shade within coffee agroecosystems reduces the spread of spores, thus providing support for the idea that shade trees within a coffee plantation act as a wind breaks and prevent local dispersal. Avelino et al. (2012) found that the surrounding landscape of pasture and/or sun coffee was highly correlated with the CLR on individual farms. Thus there is an emereging appreciation in the liteature that the transmission of spores and the viability of spores are two different forces that need to be simultaneously manage, and there exists a potential tade-off with shade between the local scale of pathogen transmission and probability of the pathogen establishing infection.

While shade may offer a way to locally reduce the spatial scale at which the pathogens can spread, we show that the final realization of the pathogen dynamics will also depend on the planting geometry of the agroecosystem. Our simulations clearly show that lattice-like planting geometries are far more susceptible to epidemic outbreaks with small changes in the spatial scale of pathogen transmission. We show that the more spatially disordered arrangement may result in higher number of plants infected at small spatial scales, but will buffer the system from epidemic thresholds of the pathogen.

While the question of what initially caused the outbreak of CLR in 2013 in Latin America is still unclear, it set up the necessary conditions to overhaul many coffee agroecosystems throughout the region. Due to the prevalence of plant death from the epidemic itself, in conjunction with the promotion of resistant varieties, most farmers throughout Latin America are replanting whole farms at a time now. This particularly important moment in the dynamics of the CLR in Latin America, as following classical agronomical recommendations for combating the CLR would mean a reduction in shade, thus increasing the scale of CLR transmission locally, as well as replanting with resistant varieties, will likely lead to planting geometries that are more lattice-like when the whole system is replanted. Studies have found that this process is already underway in parts of Central America (Valencia et al. 2018). As our study shows, the combination of shade reduction and moving back towards a uniform planting structure, increases the likelihood of the critical transition-like epidemic dynamics observed in our model.

## Acknowledgements

We would like to thank Gustavo Lopez-Bautista for help with setting up the plots to measure the empirical spatial distributions of coffee plants. Wardell Gray and Ferdinand LaMothe both provided useful feedback on the models. This work supported by NSF grant number DEB – 1853261

